# Orientation Single Molecule Localization Microscopy (oSMLM) for Decoding Orientation of Single Molecule

**DOI:** 10.1101/2023.06.27.546198

**Authors:** Prakash Joshi, Partha P. Mondal

**Affiliations:** Mondal Lab, Department of Instrumentation and Applied Physics, Indian Institute of Science, Bangalore 560012, INDIA

## Abstract

Standard SMLM facilitates the reconstruction of super-resolution map (both location and localization precision) of the target single molecules. In fact, single molecule data does provide information related to the orientation of single molecules, which can be derived from the knowledge of PSF shape and its direction. This information is vital to probe the sub-domain of macromolecules that undergo orientation and conformational changes and provides essential clues on their catalytic activity. Accessing this information in real-time opens up a powerful new window to look into the link between the orientation of macromolecules and the output function. Here, we decode the orientation of single molecules from the knowledge of PSF shape and its direction. The method is primarily based on field-dipole interaction and the fact that the distribution of emitted photons strongly depends on the orientation of the dipole (fluorophore) with respect to the polarization of light. Accordingly, the photon emission from the specimen and the resultant PSF distribution model is developed. Computational studies show changes in the shape and orientation of the recorded PSF (in the image / detector plane). Specifically, a set of three distinct distributions (Gaussian, bivariate-Gaussian and skewed-Gaussian) are recognized from the study, apart from a superset of all possible (a total of 16) distributions. Experiments were conducted on Dendra2-Actin and Dendra2-HA transfected cells that validate the emission model. We report a localization precision of *∼* 20 *nm* and an orientation precision of *±*5°. In addition, the distinct orientation of single molecules is noted for Actin and HA in a cell (Influenza type-A model). Further analysis suggests a preferred directional distribution of Dendra2-Actin single molecules, while Dendra2-HA molecules seem to be randomly oriented in a cluster. The availability of orientation information in SMLM without the need for additional optics adds a new feature, which can be explored to reveal the state of a single molecule (orientation and conformational changes) in cellular sub-domains / partitions. The study implies that the orientation of single molecules that has more profound implications for the functioning of macromolecules. The orientation information revealed by *oSMLM* technique gives it a wide-spread appeal and expands the reach of localization microscopy.

---

With the advent of super-resolution microscopy, it is now possible to unveil the nanoscopic world of living organisms. The processes that largely facilitate super-resolution are either based on stimulated emission depletion (STED) or single molecule blinking. Over the last decade, these techniques have tremendously advanced and present variants are able to resolve features down to sub-10 nm regime. Needless to say, these technique hold the potential to decipher biological processes both spatially and temporally.

Early 1990s saw the emergence of a new concept (called STED) that for the first time demonstrated the capability to surpass the diffraction limit [1] [2]. It wasn’t until early 2000s that the first far-field super-resolution technique is fully established [3] [4]. Then the year 2005-2006 saw the emergence two different techniques (structured illumination and single molecule localization microscopy) that surpassed resolution far beyond the classical diffraction limit [5] [6] [7] [8]. The ground work for single molecule based super-resolution was largely laid down by Moerner et al [9] [10]. These techniques largely convinced that it is possible to surpass the diffraction limit set by classical optics[11]. The success of super-resolution technique saw a huge interest and emergence of a family of techniques over the last decade. Most of these variants were inspired by applications in diverse disciplines across basic sciences [13], [14], [12], [15], [16], [17], [18] [19] [20] [21].

Super-resolution techniques specifically based on single molecule blinking have advanced our understanding of diverse biological process, both at cellular and organism level. For example, SMLM has revealed the nanoscopic world of macromolecular complexes and its functioning in a cellular environment including, apoptopic pores [40], nuclear pore complex [41], endocytic machinery [42], cytoskeletal structure [43]. This is made possible by the emergence of technique that are taylor-made for specific applications. Some of these include, SMLM such as, POSSIBLE, STED, MINFLUX, dSTORM, SIMPLE and ROSE that have shown sub-10 nm resolution [44] [47] [48] [49] [50] [51] [53]. Another class of microscopy techniques that have advanced the reach of super-resolution microscopy are hybrid systems. For example, the integration of light sheet and super-resolution have shown great promise [45]. It has enabled super-resolution imaging of human mammary MCF10A cell spheroids expressing H2B-PAmCherry [45]. Multicolor variant such as multicolor STORM imaging of large volumes has enabled ultrathin sectioning of ganglion cells to understand the nanoscale co-organization of AMPA receptors and their spatial correlation [35] [36] [37]. Techniques based on RESOLFT microscopy have shown 3D imaging of single molecules organization in a cell [38]. In another development, evanescent field based techniques (such as SMILE) has demonstrated volume imaging capabilities in a single cell [17] [18]. With the growing influence of super-resolution techniques, many more developments are expected in coming decade.

Although SMLM has enabled *∼ λ/*10 nm resolution, still a lot needs to be determined including, the orientation of single molecules, its actual size, interactions with neighbouring molecules and its temporal dynamics. In this respect, single molecule FRET has shown promise, but very little is know about the orientation of single molecules. However, there has been a handful of studies related to orientation. Recently, a new microscopy technique termed as single molecule orientation localization microscopy have successfully resolved structural heterogeneities in amyloid fibrils and lipid membranes [22] [23]. Machine learning is employed to design PSFs for three-dimensional imaging of single molecule distribution [24] [25]. Of-late, PSF engineering is a promising technique for revealing 3D position and orientation of single molecules [26] [27]. These handful studies does give information related to single molecule orientation, but are complex in instrumentation, cumbersome, and hard to interpret, thereby limiting its use to specific applications. So, techniques based on standard SMLM and without the need for additional hardware / optics is in high demand.

In this work, we propose a new technique to determine the orientation of single molecules in SMLM and classify them based on their PSF distribution. This is primarily based on the fact that interaction of single molecule dipole with polarized light field determines the shape of single molecule PSF and its orientation (in image plane), thereby giving discrete information related to the orientation (*θ*_*d*_, *ϕ*_*d*_) of dipole in the object plane. Such an information is revealed by the combination of PSF shape (Gaussian, bivariate-Gaussian and skewed-Gaissian) and its orientation in the image plane. In addition, analysis of oriented dipole in specific domain is also carried out, revealing the corresponding population statistics. Such information are critical in accessing biophysical process including protein-folding and its interaction with the immediate neighbourhood both in fixed and live cell.

## 1. RESULTS

### A. Optical Setup of oSMLM System

The schematic diagram of developed orientation single molecule localization microscopy (*oSMLM*) is shown in Fig. 1. The illumination sub-system is at the heart of *oSMLM*, with the activation achieved by linearly-polarized 405 nm light and excitation by a linearly-polarized 561 nm light. A series of standard optics (not shown) is used to expand, align and direct both the beam to the inverted microscope. Post interaction (between polarized light field with the single molecule dipoles), the fluorescence is collected by the high NA objective (1.3 NA, 100X, Olympus). Then the light is filtered by a series of filters arranged in a filter box (consists of notch and long-pass filters) and directed to the sensitive EMCCD detector (iXon897 Camera, Andor, UK) by a series of optics (mirrors, magnifier, lenses). Note that, a total magnification of 280X is used for obtaining large PSF. The data is collected at 30*Hz* and approximately 5000 frames are taken for each dataset. This is followed by analysis that involves, multi-functional fit (using, a generalized Gaussian function) to determine orientation and the shape of PSF.

**FIG. 1:**
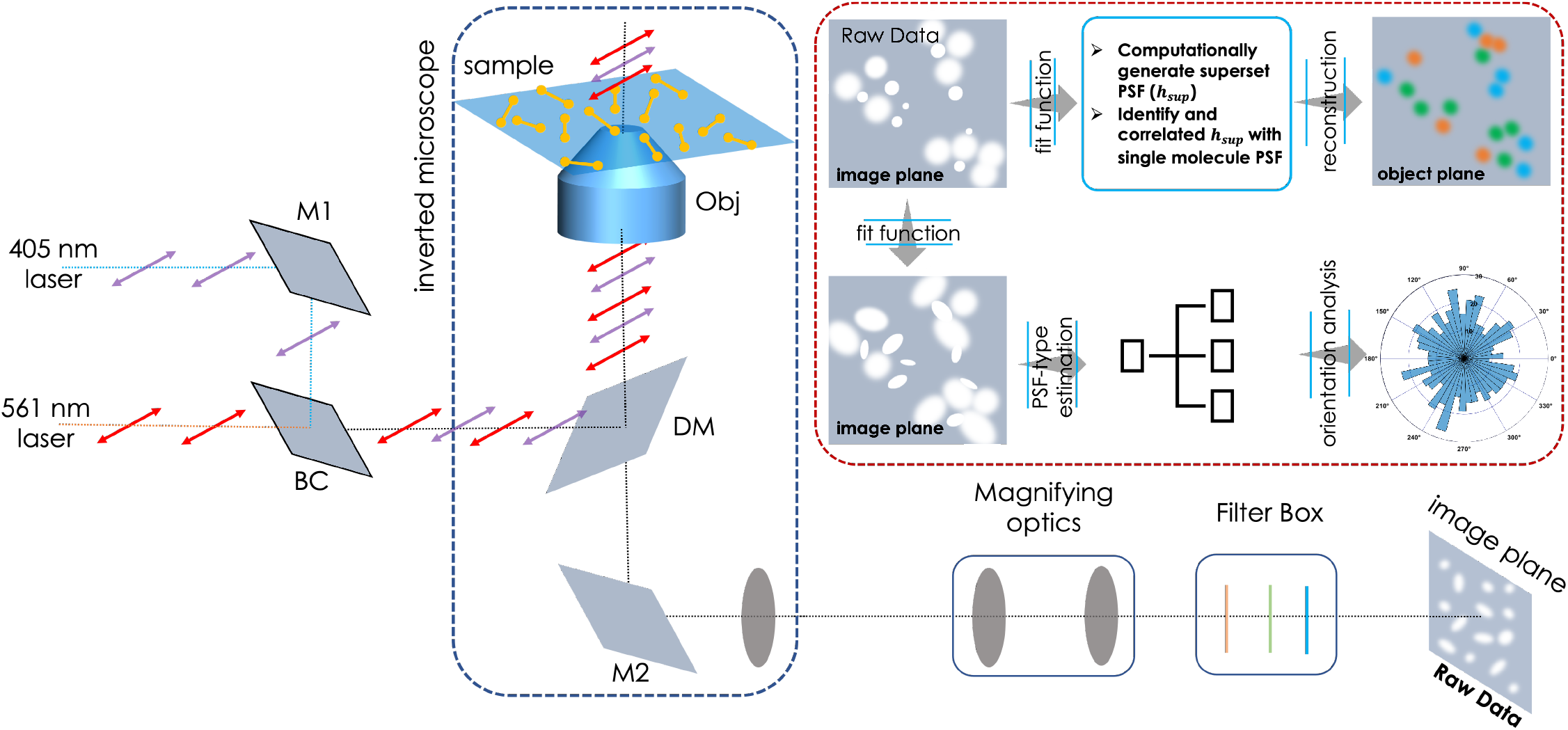
Schematic diagram of the proposed *oSMLM* system. The technique involves activation of single molecules by a linearly polarized 405 nm light and excitation by a linearly polarized 561 nm light. The interaction of polarized light with single molecule dipole determines the respective diffraction pattern (PSF) on the image / camera plane. A careful fit of the resultant PSF distribution reveals the PSF-shape classified as a set (3 PSF types i.e, Gaussian / Bivariate Gaussian / Skewed Gaussian) or a super-set (16 PSF types). In addition, *oSMLM* gives the orientation statistics of single molecules. Together these parameters determine the dipole orientation in the object plane. The inset shows further analysis for segregating single molecules based on PSF type.

### B. Characterizing Single Molecule PSF

To characterize single molecule PSF and determine its type, the detection sub-system is theoretically modelled and computational studies were carried out. Figure 2A shows the detection sub-system along with the electric field in the oil immersion medium, intermediate spaces and at the detection / image plane. Using standard classical electromagnetic theory of light (see, methods section) for a multi-layered system (see, Fig. 6), field at the focus of illumination and detection objective are given by eqn.(3), and eqn.(10), respectively. It is evident that the PSF depends on the polar angle (*θ*_*d*_) and azimuthal angle (*ϕ*_*d*_), and hence uniquely determines the orientation of single molecule dipole. The results are shown in Fig. 2B for various combination of angles, (*θ, ϕ*), with 30° ≤ *θ, ϕ* ≤ 90°. It is visually apparent that, the PSFs distribution varies for different combinations of angles. Apart from 16 PSF types, one can further simplify and classify then into three major types namely, Gaussian, bivariate-Gaussian and skewed-Gaussian (see, intensity plots in Fig. 2C). The computational study clearly indicates the role played by field-dipole interaction in determining the output PSF distribution and the dependence of the distribution on the angles (polar angle and azimuthal angle of the dipole). Based on Fig.2, we have two possible basis set for characterising single molecule PSF: Set of major PSF distribution (3 PSFs), and a super-set of all possible distributions (a total of 16 PSF distributions types).

**FIG. 2:**
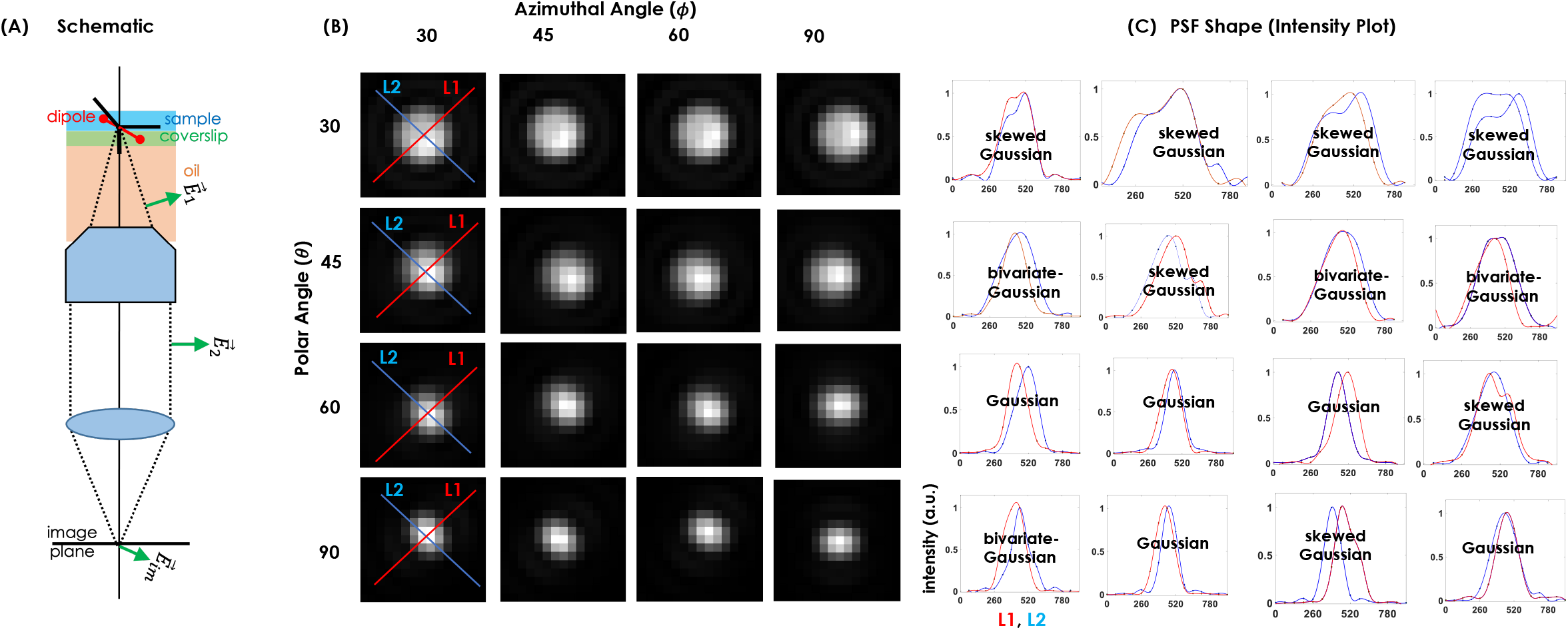
Computational study: (A) Schematic diagram showing fields at the front and back focal plane, and in the image plane. Field is calculated based on the developed model of field-molecule interaction (see, Materials and Methods section) that incorporates the effect of sample, coverslip and immersion oil refractive indices. (B) The resultant PSF for varying combination of (*θ, ϕ*); 30° ≥ *θ* ≤ 90° and 30° ≥ *ϕ* ≤ 90 °. (C) Corresponding intensity plots suggests 16 PSF types and specifically three distinct types of PSFs (Gaussian / Bivariate Gaussian / Skewed Gaussian).

To establish the theoretical findings and computational study, the *oSMLM* system is built and single molecule experiments are carried out. NIH3T3 cells were transfected with photoactivable probes (Dendra2-Actin and Dendra2-HA plasmid DNA). Subsequently, the cells were fixed for data collection following standard protocol post 24 hrs of transfection [44] [46]. Figure 3A shows a transfected cell image of which a small region is imaged. The intensities used for activation and excitation lasers are, 112 *μW* and 6.12 *mW*, respectively. The data / images are recorded at 33 *Hz* and a total of 5000 images are collected. *oSMLM* reconstructed images and the respective enlarged sections (R1, R2 and R3) show the orientation of single molecule PSFs along the actin filaments. In another study, *oSMLM* is used to understand the clustering behaviour in an influenza (type A) model. Healthy NIH3T3 cells were transfected with Dendra2-HA plasmid DNA following standard protocol [44]. The cells were fixed and used for super-resolution imaging to understand the clustering of HA molecules. Figure 3B shows both *oSMLM* reconstructed images along with the enlarged sections (R1, R2, R3) of few HA clusters. The orientation of individual HA molecules in the clusters indicate a heterogeneous mix of dipoles. The corresponding histogram plot (Fig. 3(C,D)) shows the distribution of molecules (PSF types) as Gaussian, bivariate-Gaussian and skewed-Gaussian, with bivariate appears to be the most abundant. This is a sharp departure from the information available from SMLM and variants that blindly assume 2D Gaussian which may be an over-estimation and must be corrected for accurate representation of single molecule data. In addition, Fig. 3(C,D) shows the histogram plots of the number of single molecules oriented in a specific angle. Both Dendra2-Actin and Dendra2-HA transfected cells were considered and the population statistics of two class i.e, bivariate and skewed Gaussians are determined. It is evident that, molecules with bivariate-Gaussian distribution are isotropically distributed in the cell, whereas skewed-Gaussian have preferential directions. Specifically, they are mostly directed in the range, [30° − 60°] and [22° − 60°] for actin and HA, respectively. Detailed orientation statistics is tabulated in Table 1. The orientation of a molecule is an important indicator of conformational changes and catalytic activity. The availability of such discrete information at single molecule level is seldom possible with the existing SMLM microscopy techniques, and so *oSMLM* may play a major role in understanding interactions and other processes (such as conformational changes and protein folding) at single molecule level.

**TABLE I:** Orientation Statistics for Dendra2-Actin and Dendra2-HA single-molecule proteins in a transfected NIH3T3 cells.

| Protein | Photon Distribution | Orientation (%) | | | $lp_x$ (nm) | $lp_y$ (nm) | $lp$ (nm) |
| --- | --- | --- | --- | --- | --- | --- | --- |
|  |  | 0 – 60° | 60 – 120° | 120 – 180° |  |  |  |
| Dendra2-Actin | Gaussian | – | – | – | – | – | NA |
|  | Bivariate Gaussian | 37 | 32 | 31 | 16 | 18 | – |
|  | Skewed Gaussian | 38 | 30 | 31 | 18.6 | 18 | – |
| Dendra2-HA | Gaussian | – | – | – | – | – | NA |
|  | Bivariate Gaussian | 41 | 30 | 26 | 24.3 | 29 | – |
|  | Skewed Gaussian | 40 | 33 | 27 | 21 | 19 | – |

**FIG. 3:**
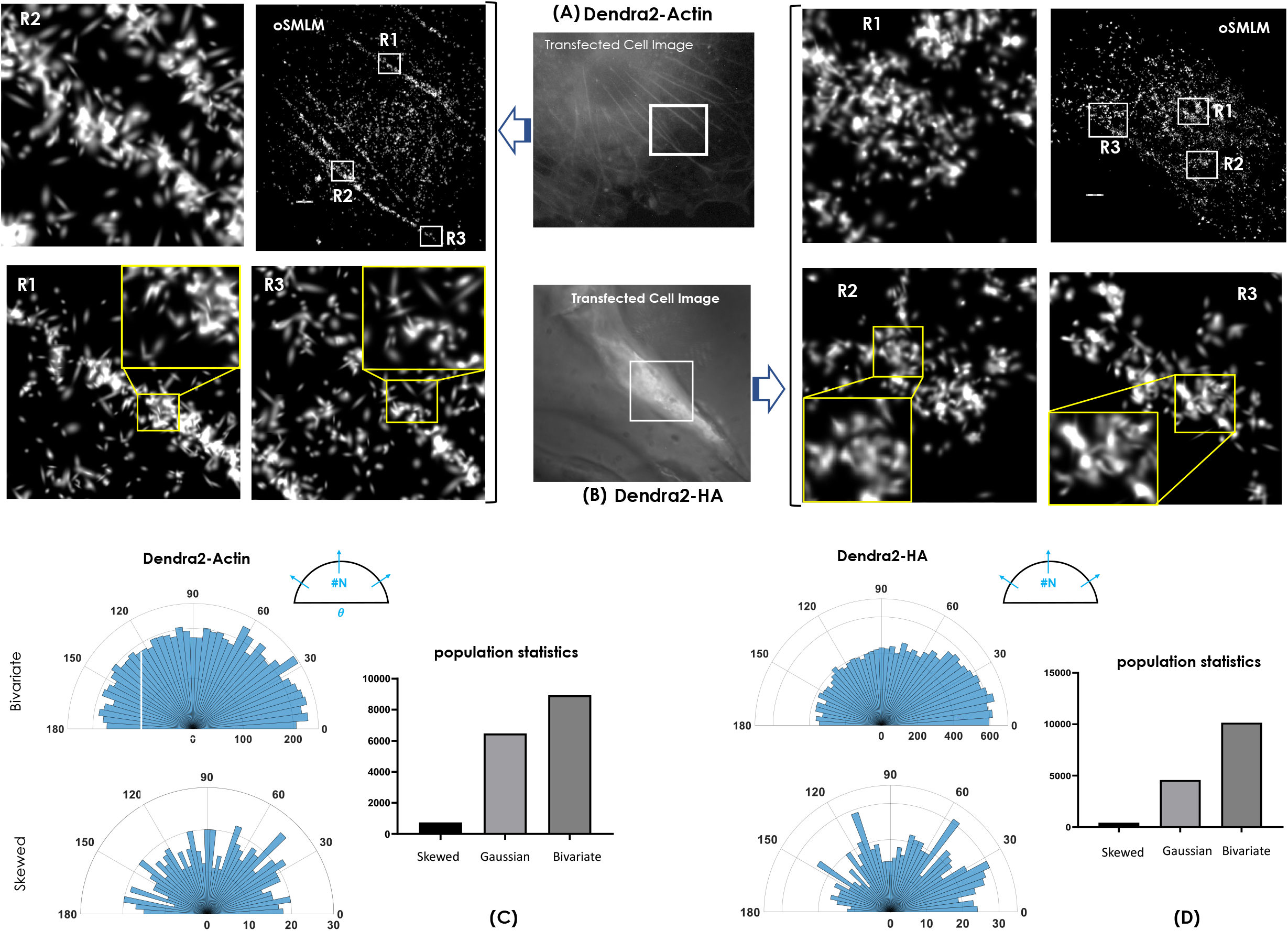
Modelled photon distribution of recorded single molecule PSFs. (A) Reconstructed map of Dendra2-Actin molecules in the image plane. Enlarged image of three sub-regions are also displayed, which shows the orientation and PSF shape of individual single molecules. (B) Reconstructed map of Dendra2-HA molecules in the image plane, suggesting clustering of HA molecules in a transfected NIH3T3 cell. However, *oSMLM* gives additional information related to the orientation and PSF type of individual single molecules. (C,D)Orientation statistics of bivariate and skewed Gaussian PSF for both Dendra2-HA and Dendra2-Actin along with the population statistics. Scalebar = 1*μm*.

In addition to standard parameters (centroid, number of photons and localization precision), the data provides multi-class perspective (in terms of orientation) of single molecule. Figure 4 displays multi-class data representing the abundance and arrangement of different PSF types (Gaussian, bivariate-Gaussian and skewed-Gaussian) of single molecules in image place. Here, specific colors are used to represent PSF type. Few enlarged regions are also shown that clearly indicate the abundance of specific class of molecules. Statistical analysis of single molecule population indicate the dominant presence of Bivariate as opposed to Gaussian. This is a departure from the general assumption made in existing SMLM techniques forcing all the single molecules to have a Gaussian distribution. In addition, we did observe a small non-negligible fraction of molecules with skewed-Gaussian distribution. In order to understand the impact of polarized light field on single molecule, we have also characterized localization precision. The corresponding statistics (*l*_*x*_ and *l*_*y*_) in the lateral plane is displayed in Fig. 4. This clearly indicates distinct distribution for Dendra2-Actin and Dendra2-HA suggesting a broad spectrum of localizations along X and Y i.e, *l*_*x*_ and *l*_*y*_. It is quite evident from the respective plots (Fig.4 C and D) that the localizations are not symmetric, thereby contradicts the assumption of Gaussian photon distribution in the presence of polarized light field. In addition, the localization plot suggests localization along x (*l*_*x*_) seems to suffer more compared to *l*_*y*_ in the case of for Dendra2-Actin. The difference-plot (Fig. 4C,D insert) suggest a large difference between *l*_*x*_ and *l*_*y*_, for Dendra2-Actin and Dendra2-HA, respectively. Overall, interaction of single molecules for a specific state of polarization (along X) brings-in non-symmetrical photon distribution, giving information on the orientation and immediate neighbourhood of single molecule.

**FIG. 4:**
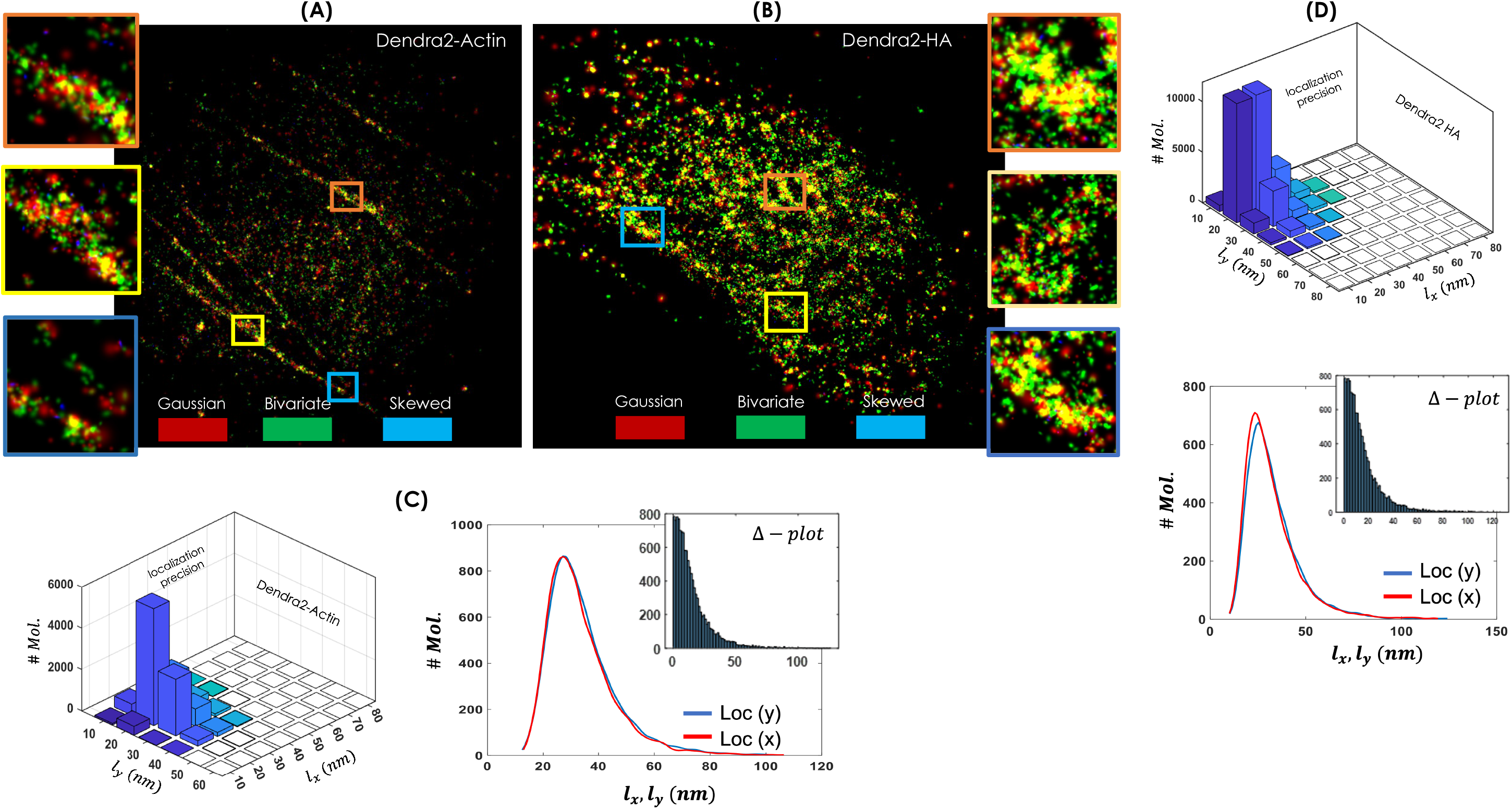
Modeled PSF-type Encoded Map of the Recorded Single molecule PSF in the image plane using Set (Gaussian, bivariate-Gaussian, skewed-Gaussian): (A) Pseudo-colored image of single molecules in Dendra2-Actin transfected cell where, the color indicate the PSF-type. Alongside, the localization precision *l*_*x*_ and *l*_*y*_ for single molecules (both bivariate and skewed Gaussian) is also shown. (B) Pseudo-colored map of single molecules in Dendra2-HA transfected cell along with the localization map. Three separate enlarged sections are also shown that suggest a mix of single molecule types. This shows the arrangement and orientation of single molecules along the actin filaments and in a HA cluster. (C,D) Localization precision plots along X and Y for (A) and (B), respectively. Alongside, difference plots (Δ-plots) are also shown for both the cases.

### C. Orientation Encoded Reconstructed Map

Finally, superresolution image (in object plane) is reconstructed along with the orientation information inscribed on it (see, Fig. 5A,B). To do so, we have computationally determined the PSF based on the orientation angles (both azimuthal and polar angles). Theoretical modelling of photon emission along with the beam propagation in the detection system is carried out as discussed in the Material and Methods section (Theory and Computational Study). Specifically, eqn.(8-10) are computationally implemented and the system PSFs are determined for different values of orientation angles (30 ≥ *θ, ϕ* ≤ 90). The study is carried out on a grid size of 15 *×*15 pixels with a *XY* sampling of 65nm. The combination of *θ* and *ϕ* angles give rise to a set of 16 possible PSF types which can be considered as the basis set for recognizing the orientation angles of the single molecule dipoles (fluorophores). The next step involves the segregation of recorded single molecules into these 16 PSF basis-sets (super-set). This is accomplished by correlating the experimentally recorded single molecule with the basis-set PSFs (see, Fig. 5C). Correlation factor, *η >* 0.92 is considered and the corresponding single molecule PSF is segregated into super-set PSF types and a distinct color is used to represent them. So, the single molecule PSFs / dipoles are color-coded based on their orientation (combination of azimuthal and polar angles) in the object space. Analysis reveals the orientation statistics of dipoles for both the cases (cells transfected with Dendra2-Actin / Dendra2-HA) as shown in Fig. 5D. Specifically, we noted orientation of dipoles with azimuthal and polar angle (*θ, ϕ*) at 90° is dominant for Dendra2-Actin, whereas, clear dominance of dipoles at, (*θ* = *ϕ* = 30°) is evident for Dendra2-HA. This gives insight into the orientation preference of single molecules while performing complex biological processes (such as, clustering) that has deeper biological significance.

**FIG. 5:**
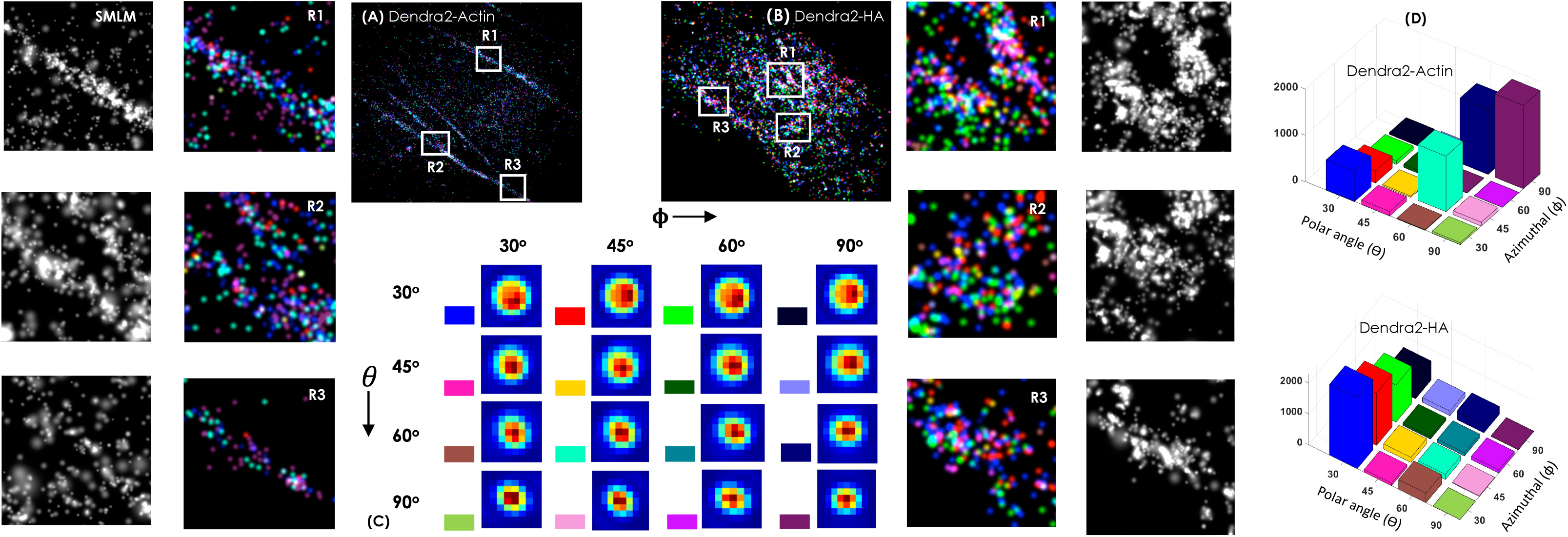
Reconstructed image (encoded with orientation information) in the object plane Using Superset (16 PSF types): (A) Orientation (*θ, ϕ*)-encoded super-resolved image of Denra2-Actin transfected image. Few enlarged sections (R1, R2, R3) are shown that imdicate the dominance of small orientation angles (30 ≤ *θ, ϕ* ≤ 45) (light-blue) as compared to large angles (60 ≥ *θ, ϕ ≤* 90). (B) Orientation (*θ, ϕ*)-encoded super-resolved image of Denra2-HA transfected image. Few enlarged sections are shown indicating more dominance of red as compared to blue color. Alongside SMLM reconstructed images are also shown. (C) The basis superset is represented by different colors for a combination of (*θ, ϕ*). (D) Population statistics of single molecules (Dendra2-Actin and Dendra2-HA) at different combinations of polar and azimuthal angles.

**FIG. 6:**
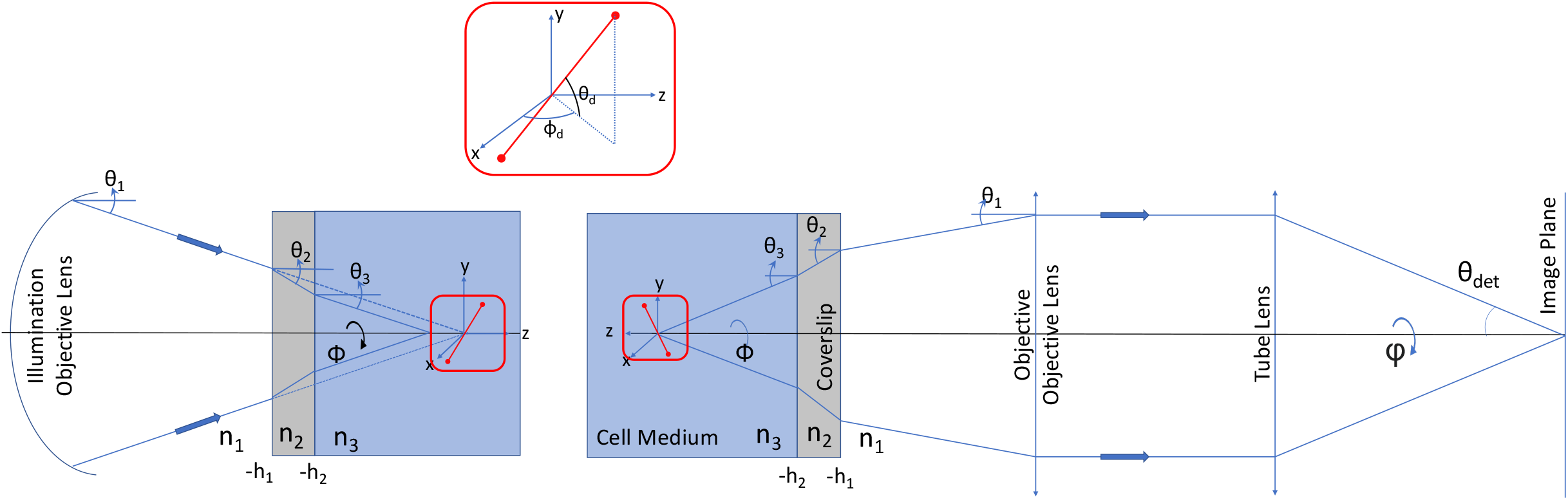
Electromagnetic wave traversing through multiple mediums in a lens focussing system. The diagram shows propagation of electromagnetic field through multiple interfaces and its interaction with the single molecule dipole (red color) in an *oSMLM* imaging system. All the parameters necessary to model the field (both in the illumination and detection sub-systems) in the complex imaging system are shown. Accordingly, the fields are computed and image of the radiating dipole is formed.

## Conclusion & Discussion

A new localization microscopy technique (*oSMLM*) is developed for decode the orientation of single molecules. The technique use polarized light field to excite dipole (single molecule) and the resultant field-dipole interaction gives rise to unique photon distribution pattern (PSF) for single molecule (in the image / camera plane), revealing information related to its orientation. Based on the study, we noted that there are three dominant types of single molecules PSFs (Gaussian, bivariate-Gaussian, skewed-Gaussian), although the complete superset comprises of 16 PSF types. These details at the single molecule level gives a plethora of information about the location, orientation, and overall state of the molecule.

Two beams (activation and excitation beams), both linearly-polarized are used to illuminate the single molecule dipoles, and the resultant photon distribution of single molecules (PSF) is recorded by a 4f-widefield detection system. Unlike existing SMLM techniques that use 2D Gaussian function as a preferred fit for the detected spots, the proposed *oSMLM* assumes a generalized-Gaussian function. The advantage of using this function eliminates the need for forcing a pre-assumed function and lies in its flexibility to use non-symmetric PSF distribution. The recorded data is fit and the relevant parameters (related to uniformity and skewness) are estimated. This is then used to determined separate localization precisions (*l*_*x*_ and *l*_*y*_) and orientation (*θ*_*d*_, *ϕ*_*d*_) for further processing. To validate, theoretical model is conceived and computational studies are carried out that resonate with a physical system mimicking field-molecule interaction in the specimen (see, Fig. 2 and II.A). Computational study clearly demonstrate the effect of field polarization (along 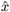) on the dipole orientation (*ϕ*_*d*_, *θ*_*d*_). A careful analysis of the PSF (using intensity plots) reveal three major distinct class of single molecules, closely resembling normal-Gaussian, bivariate-Gaussian and skewed-Gaussian for 30 ≥ *θ, ϕ* ≥ 90. This directly links the type of dipole orientation in the object plane. Although, a much more complete PSF characterization is revealed by the computational study which is a super-set of 16 possible PSF types (see, Fig. 2 and Fig. 5). So, in-principle, the decoded orientation information of the single molecule PSF can be incorporated in the reconstructed process of super-resolution map. The simulation results have provided a firm ground for further investigation and experimental validation.

Single molecule signatures are recorded with the *oSMLM* optical set-up and processed. Two samples are considered, first NIH3T3 cells transfected with Dendra2-Actin for visualizing the organization of single molecules on a Actin filament, and the second sample is cells transfected with Dendra2-HA for visualizing HA molecules distribution in a cluster. *oSMLM* revealed the orientation information of the molecules in an organelle (Actin filaments) and in a HA cluster (see, Fig. 3). Finally, the recorded single molecule PSFs are segregated into different types and statistical analysis were carried out to visualize the dominant type (see, Fig. 4). Alongside the localization precision in the lateral plane (both along X and Y) are carried out to access the PSF shape information. This suggests that a large number of molecules are better localized along *y* than *x*-direction, indicating a departure from the notion of isotropic localization which is a prevalent assumption in SMLM and most of its variants. Using these information related to the orientation of a molecule, its position (centroid) and localization precision, we were able to reconstruct a super-resolution map with orientation information engraved on it (see, Fig. 5). Specifically, a set of 16 possible orientations (a combination of polar and azimuthal angles (*θ*_*d*_, *ϕ*_*d*_)) of the single molecule dipole is constructed. An important consequence of this is to determine the preferential orientation of single molecules in different biological processes (single molecule clustering in Influenza pathogenesis or its location on organelles such as, Actin filaments). In addition, statistical analysis and population studies are carried out to determine the dominance class of molecules for both the cases, showing preferential orientation states. Such informations are valuable in cell and disease biology to understand the role of single molecule orientation in critical biophysical processes such as, protein-folding, conformational changes.

To our knowledge, this is the first time orientation information is revealed from a SMLM data collection, without the need of additional sophisticated optics. Indeed, single molecule data is rich enough to reveal its orientation and the consequences (related to single molecule neighbourhood, and folding dynamics) arising out of it. This goes on to show that, *oSMLM* advances localization microscopy by adding an important feature thereby expanding the reach of single molecule super-resolution microscopy.

## II. MATERIALS AND METHODS

### A. Theory

#### Illumination PSF

Consider light is being focussed by an aplanatic lens propagating through multiple refractive index mediums (oil, coverslip glass and specimen of refractive index, *n*_1_, …, *n*_*N*_, respectively) as shown in Fig. 6. The incident light is assumed to be linearly polarized along *x*-axis. Following Torok et al. [28], the resultant electric field at the geometrical focus of the illumination system is given by,

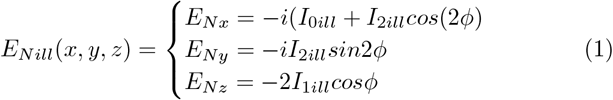

where,

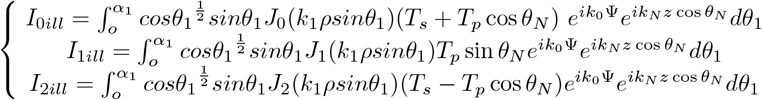

where, the abberation function is, Ψ_*I*_ = *h*_*N−*1_*n*_*N*_ cos *θ*_*N −*_ *h*_1_*n*_1_ cos *θ*_1_. Here, *z* = *h*_1_, *h*_2_, …, *h*_*N*_ represent interfaces as shown in Fig. 6.

The components of polarization vector are defined as,

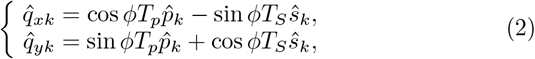

which is a function of basis polarization vectors 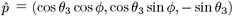 and *ŝ* = (−sin *ϕ*, cos *ϕ*, 0) and depends on the Fresnel coefficients *T*_*a*_ = *T*_*a*,3*−*2_, *T*_*a*,2*−*1_ for 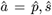. The Fresnel coefficients for the two contributing interfaces are defined by, *T*_*a*,1*−*2_ = 2*c*_*a*,1_*/*(*c*_*a*,1_ + *c*_*a*,2_), with *c*_*p,l*_ = *n*_*l*_*/cosθ*_*l*_ and *c*_*s,l*_ = *n*_*l*_ cos *θ*_*l*_ for *l* = 3, 2, 1. Note that, spherical coordinates system (with, 0 ≤ *θ* ≤ *π* and 0 ≤ *ϕ* ≤ *π*) is used for representation.

The resultant electric field and the illumination PSF are given by,

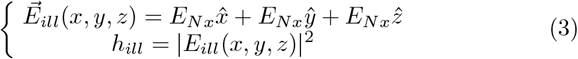

At the geometrical focus, the field interacts with the fluorescent molecules, which act as electric dipoles in the presence of incident electric field 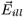. This results in the excitation and subsequent emission of light (fluorescence) from the radiating dipole (fluorescent molecule).

#### Detection PSF

##### Case I: Oil Immersion Objective

Post interaction, the dipole radiates isotropically and a part of which is collected by the detection objective. The emitting dipole is assumed oriented in a direction given by the unit vector, 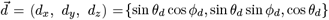, where *θ*_*d*_ and *ϕ*_*d*_ are respectively the polar and azimuthal angles (see, Fig. 6). The electric field of the radiation field is given by,

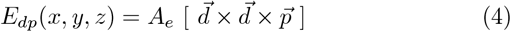

where, *A*_*e*_ is the constants of proportionality, 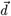 is a unit vector along the dipole, and 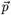 is the electric dipole moment.

Considering the realistic situation that incorporates the effect due to immersion oil, cover slip and specimen. The electric field 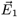 at the front focal plane of the objective is can be directly written following Torok et al., [29] i.e,

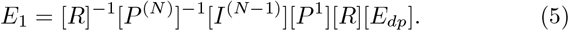

where, the metrics, [*R*] is a rotational matrix corresponding to coordinate transformation, [*P*] is the rotational matrix for rotated *s* and *p* components of electric, and [*I*] corresponds to the change in refractive indices at the interface, with *N* as the number of mediums involved.

Following Ref. [29] [30], the explicit form of the matrices are given by [28][29],

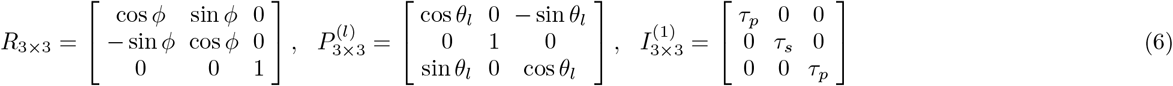

where, *τ*_*p*_ and *τ*_*s*_ are the Fresnel coefficients given by, 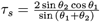, 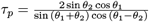 [28].

The objective lens collimates the field, 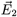 which can be obtained from 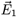, by noting the coordinate transformation from spherical-polar to cylindrical system. Incorporating the effect of lens, the field 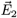 becomes [30],

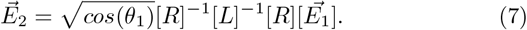

where, 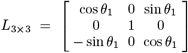 describes the changes in the electric field as light propagates through the objective lens.

Finally, the electric field in the image plane can be obtained by taking Fourier transform of 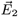, component-wise. So, the field components in the image plane are [31],

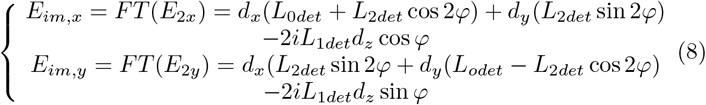

where,

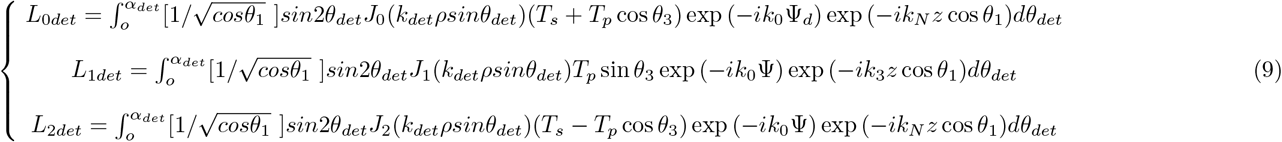

and, Ψ_*D*_ = *h*_1_*n*_1_ cos *θ*_1_ *− h*_*N−*1_*n*_*N*_ cos *θ*_*N*_.

Here, *α*_*det*_ and *k*_*det*_ are the angular aperture angle of detector lens and wavenumber in the image plane, respectively. The relation between *α*_1_ and *α*_*d*_ is given by,

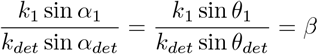

where, *β* is the magnification of the detection system.

The corresponding detection PSF (Photon distribution) can be obtained from the electric field in the image plane i.e,

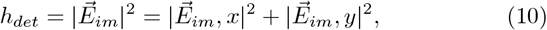

##### Case II: Air Objective

Employing air objective for detection is one of the easiest implementation and thus worth considering. In case of air objective, the electric field *E*_1_ at the front focal plane of the objective lens (air objective lens) is given by the classical relation by Born and Wolf [32]. Since there is no change of refractive index the dipole field (*E*_*dp*_) and field at the front focal plane (*E*_1_) are the same i.e, *E*_1_ = *E*_*dp*_. The field in image plane is given by,

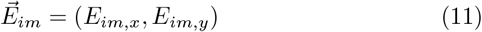

and the diffraction integrals (*I*_0_, *I*_1_ and *I*_2_) are as derived in Boivin and Wolf formulation [32].

The corresponding field components are [31],

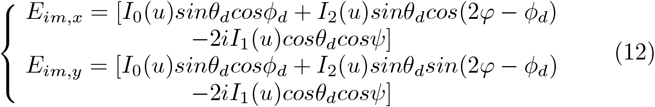

where, the amplitude is taken as unity and the diffraction integrals are given by,

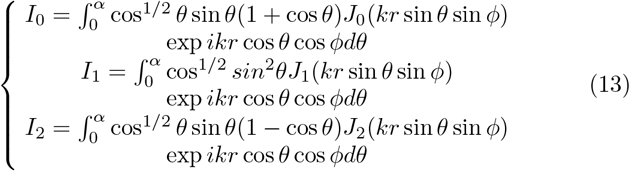

### B. Computational Study

Computational studies were carried out using eqn.(10) to understand the effect of field polarization on single molecule dipole [33] [31]. Based on the present optical configuration and sample preparation, the refractive indices for immersion oil, coverslip and the specimen are taken as, 1.52, 1.52, and 1.33, respectively. The study was carried out on a grid size of 15 *×* 15*μm*^2^ with a sampling of 60nm in the lateral *XY* -plane. The results are shown in Fig. 2, and discussed in the Results section.

### C. Correlation Study

For correlation study MATLAB inbuilt function “corr2” is used. The purpose of this study was to analyse the degree of similarities between the simulated data at the object plane and the image plane. To correlated the image, we have considered simulated data (superset) as the basis set. The corr2 function calculates the correlation coefficient, which defines the linear relationship between the pixel values of the two images (PSF image and raw image containing single molecules). The correlation coefficient of +1 indicates a complete match of the two images, 0 implies no correlation. In our study we have chosen a cutoff of 0.92 as the match parameter, which indicates anything greater than 0.92 is considered as matched. The following expression is used for calculating correlation coefficient,

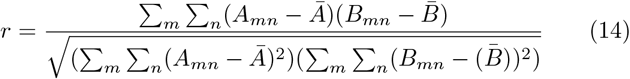

where *Ā* and 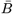 are the mean of A and B respectively.

For computational study, a dipole polar and azimuthal angles of, 30, 45, 60, 90 degrees were considered, and the PSF was generated. on a grid of 15 *×* 15 pixels. The raw data image was reconstructed from the parameter extracted from *nlinfit* of the raw data using the generalised Gaussian equations. The raw data image was correlated with the basis set and if the correlation coefficient is found greater than 0.92 then it is noted and color coded with a color. Each combination of polar and azimuthal angle is represented by a different color (distinguished color) and they are shown in the Fig. 5.

### D. Cell Sample Preparation

NIH 3T3 Cells are thawn from -80 degree and then centrifuged in equal amount of media to remove DMSO content from the cells. The palette of cells are then resuspended again in 1ml of culture media (DMEM, FBS and antibiotics) and seeded in the T25 flask with a count of 1lakh to 2lakh cells using hemocytometre, and kept in incubator (5% *CO*_2_ and 37°C)for 3 days to get a confluency of 80-90%. The confluent cells are then trypsinated and splitted two to three times (two to three passage) before going for the experiment.

### E. Transfection Protocol

NIH 3T3 cells were trypsinated from t25 flask and centrifused to get the palatte. These palette were resuspended in culture media (1ml). The cells were counted by hemocytometre and 1lakh cells were seeded in bottom coverslip 35mm dish or with the coverslip inserted in 35mm dish. For the later case the coverslip was washed with PBS twice and kept in UV for 1 hr and then it is washed with ethanol and dried in the laminar flow and then the coverslip is inserted in new 35 mm dish. We have put a small amount of culture media on the coverslip (to make it coated with high nutrient) and then kept it in incubator to let the media dry so that the coverslip act as a coated coverslip. Then we have put 1.5 ml of culture media on 35 mm dish on top of that we have put 1 lakh cells and were shaked properly so that the cells are spreaded all through the coverslip. Then the cells were transfected with D2HA and D2Actin plasmid after 12 -14 hr of seeding. We have used the manufacturer protocol of Lipofectamin 3000. The cells were kept in incubator for 15 min after putting lipo-plasmid complex without any other media. After 15 min 500 ul of optimum is added to 35 mm dish and kept for 3 hr in incubator. Post 3 hr DMEM(500ul,without FBS) is added and then kept for another 16-18 hr. The cells were then incubated for 36 hr more with the addition of 700ul of culture media. The cells were washed with PBS 3 times and then incubated with paraformaldehyde for 15 min at room temperature and then washed again with PBS. After fixing, the cells were sealed in the glass slide using Fluorosave solvent (Invitrogen, Carlsbad,CA, USA) to preserve for a long duration. The glass slide is then observed in the microscopy and the experiments were carried out, both in confocal and super resolution microscopy.

### F. Data Analysis

The image is captured by EMCCD camera which is operated at an exposure time of 33ms. The recorded data is then processed to identify and localize single molecules. The process begins with background subtraction using rolling ball algorithm in MATLAB and then the brightest spots are identified. Note that, if two bright spots are closer than 1.5 times of the diffraction limit, both the molecules are discarded. The identified bright spots are then fitted with a generalised gaussian *f*_*p*_ using non-linear square fitting algorithm [34],

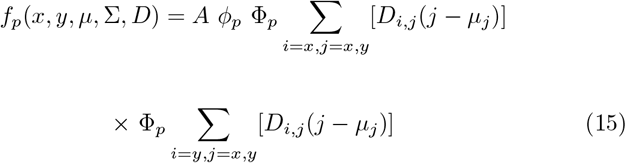

where, *ϕ*_*p*_(*x, y, μ*, Σ) and Φ_*p*_(*x, y, μ*, Σ) denote the probability density function and cummulative distribution function of p-dimensional normal distribution (with mean *μ* and covariance matrix Σ) given by [34],

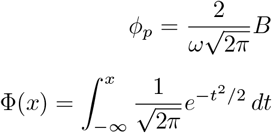

where, *B* = *A*_1_ exp (*−*(*a* * (*X − x*_0_).^2^ + 2 * *b* * (*X − x*_0_). *\** (*Y − y*_0_)+*c*\* (*Y − y*_0_)^2^)), and,

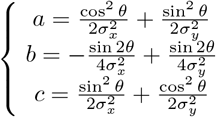

Here *D* is defined as the shape of the Gaussian with, (*x*_0_, *y*_0_) as the centroid and *ω* as the full width half maxima of the Gaussian,

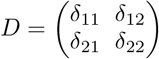

The fitting function facilitates the determination of the centroid of single molecule, its FWHM along x and y, its shape shape, and the angle of orientation. Subsequently, the localization precision is calculated using the following expression,

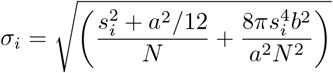

where, *i* = *x, y*. Here, *N* is the number of collected photons, *a* is the pixel size of the detector, *b*^2^ is the average background signal and *s*_*i*_ is the standard deviation of the PSF.

## Acknowledgements

The authors acknowledge financial support from parent institute (Indian Institute of Science, Bangalore, India). PPM conceived the idea. PJ, and PPM carried out the experiment. PJ prepared the samples. PPM wrote the paper by taking inputs from PJ.

## Data Availability

The data that support the findings of this study are available from the corresponding author upon request.

